# Capture of circulating metastatic cancer cell clusters from a lung cancer patient can reveal a unique genomic profile and potential anti-metastatic molecular targets: A proof of concept study

**DOI:** 10.1101/2023.09.19.558270

**Authors:** Kourosh Kouhmareh, Erika Martin, Darren Finlay, Anukriti Bhadada, Hector Hernandez-Vargas, Francisco Downey, Jeffrey K. Allen, Peter Teriete

## Abstract

Metastasis remains the leading cause of cancer deaths worldwide and lung cancer, known for its highly metastatic progression, remains among the most lethal of malignancies. The heterogeneous genomic profile of lung cancer metastases is often unknown. Since different metastatic events can selectively spread to multiple organs, strongly suggests more studies are needed to understand and target these different pathways. Unfortunately, access to the primary driver of metastases, the metastatic cancer cell clusters (MCCCs), remains difficult and limited. These metastatic clusters have been shown to be 100-fold more tumorigenic than individual cancer cells. Capturing and characterizing MCCCs is a key limiting factor in efforts to help treat and ultimately prevent cancer metastasis. Elucidating differentially regulated biological pathways in MCCCs will help uncover new therapeutic drug targets to help combat cancer metastases. We demonstrate a novel, proof of principle technology, to capture MCCCs directly from patients’ whole blood. Our platform can be readily tuned for different solid tumor types by combining a biomimicry-based margination effect coupled with immunoaffinity to isolate MCCCs. Adopting a selective capture approach based on overexpressed CD44 in MCCCs provides a methodology that preferentially isolates them from whole blood. Furthermore, we demonstrate a high capture efficiency of more than 90% when spiking MCCC-like model cell clusters into whole blood. Characterization of the captured MCCCs from lung cancer patients by immunofluorescence staining and genomic analyses, suggests highly differential morphologies and genomic profiles., This study lays the foundation to identify potential drug targets thus unlocking a new area of anti-metastatic therapeutics.

## Introduction

### Significance of metastatic cancer cell clusters

Metastasis remains the primary cause of cancer-related deaths. A recently published clinical analysis identifies metastasis as the cause of death in 50.2% to 90.4% of cases, depending on the origin of the primary tumor. The resulting high metastatic death rates indicate that increased focus on anti-metastatic therapy is urgently needed. Many cancer types have a demonstrated propensity to establish distant metastases among a defined set of organs.

Non-small cell lung cancer (NSCLC), a major malignancy with high mortality, typically metastasizes to bone, followed by brain, liver and adrenal glands.[1] It is a prime example of mortality driven by distant metastases to many vital organs.[2, 3] However, the mechanism driving the spread, in particular cancer cell transportation and immune evasion is not well understood. The inability to isolate and profile circulating metastatic cells constitutes a major challenge in understanding these pathways.

Over the past decade, tremendous strides in developing better therapies via molecularly targeted and immune-checkpoint inhibitor treatments, have improved patient outcomes.[4] However, similar significant gains have not been observed for metastatic disease.[5] A key factor in this discrepancy is the complexity of cancer metastasis. Evidence has shown that signaling pathways driving metastatic lesions are often different than found within the primary tumor.[6] A recent publication from Anderson *et al.*, representing the Cancer Research UK and Australia Metastasis Working Group stated: “The standard cancer drug discovery and development pathway, including that for molecularly targeted and immunotherapies, generally ignores the ability of experimental medicines to inhibit metastasis.” The authors went on to highlight: “To treat metastasis effectively, we must inhibit fundamental metastatic processes and develop specific preclinical and clinical strategies that do not rely on primary tumor responses.”[7]

Extensive research into individual circulating tumor cells (CTCs), during this past decade, has already begun to shed light on tumor proliferation but the underlying mechanism of metastasis remains largely unknown.[8] Adding further complexity in understanding these mechanisms, the majority of individual cancer cells in circulation undergo anoikis or are eliminated by the immune system.[9] However, an important smaller subset of CTCs circulates in the bloodstream as a cluster of two or more cells. Compelling evidence has demonstrated that CTC clusters have a significantly higher tumorigenic potential once they enter circulation as opposed to individual CTCs.[10–12]

Regarding circulating clusters or metastatic clusters, publications have referred to them with different nomenclature such as: circulating tumor cell clusters (CTC clusters),[13] circulating tumor micro-emboli (CTMs),[14] clustered circulating cancer stem cells (cCSCs),[15] metastatic cancer cell clusters (MCCCs),[16] or by using a more general term called collective cell migration,[17] yet all share common characteristics such as a being composed of 2-50 cells with up to 100-fold higher metastatic potential than individual CTCs.[13, 18] [19] Here we will use the terms MCCCs and CTC clusters interchangeably to refer to these circulating, highly metastatic entities.

A number of recent studies have begun to elucidate the phenotypic composition of these MCCCs and have shown them as either homotypic, composed of all cancer cells, or heterotypic, containing multiple cell types including non-cancerous cells.[20–22] The MCCC phenotype possesses an adaptive mechanism that enhances their survival in the harsh bloodstream environment contributing to their metastatic potential.[23–26] Specifically, heterotypic MCCCs demonstrate an increased ability to evade immune surveillance which is postulated to be a result of cancer cell and immune cell communication within the cluster.[27–29] The metastatic cancer cell clustering phenomenon is also thought to mitigate an immune response from natural killer (NK) cells, due to surface receptors in addition to intercellular communication present within CTC clusters.[30, 31]

New drug therapies, focused on MCCCs, will have the potential to dramatically reduce metastasis and improve patient survival. A landmark paper published in 2019 by Aceto *et al.* demonstrated that targeting MCCCs significantly reduces metastasis in an orthotopic breast cancer model in mice.[32] Genomic analysis of breast cancer clusters from patients revealed 10 differentially regulated pathways unique to metastatic clusters.[32] Cardiac glycosides, an existing class of drugs targeting one of those pathways, was evaluated in an *in vivo* animal model. Treatment with one of the drugs (Ouabain) reduced the number of CTC clusters by 60% and decreased metastasis to the lungs and other organs by 80-fold. An intriguing aspect from this work is that the Ouabain dosage levels utilized are not cytotoxic and only disrupt the MCCCs leading to a reduced metastatic index *in vivo*. This study strongly supports the necessity to characterize the MCCCs from patients, to identify previously unseen therapeutic targets and ultimately develop novel drugs thus opening a completely new paradigm in anti-metastatic therapy.

### Current Tumor Cell Isolation Methods from Whole Blood

Insights into how solid tumors migrate to specific pre-metastatic sites is necessary to develop focused therapeutic approaches that can prevent metastasis. While the importance of MCCCs has been well established, technologies to capture and characterize the MCCCs remain problematic and has limited research into this field.

Existing technologies used to capture CTC clusters or MCCCs, rely on systems designed to capture single CTCs. Many technologies are based on a size filtration mechanism for capture, relying upon the premise that individual CTCs are larger than most leukocytes.[33, 34] Using size filtration, patient blood samples are processed and then sorted based on the individual cell diameter. Whole blood consists of three main components: platelets, red blood cells, and white blood cells. White blood cells, which are the largest of the three components, are generally 12 - 15µm in size. The larger MCCCs, when present in whole blood, would be isolated and available for further profiling. Although the main benefit of size-based technology is that its label-free and requires little pre-processing, it has major limitations when it comes to successfully identifying and isolating several types of individual CTCs and MCCCs. It is possible for smaller MCCC’s and many CTC’s to be falsely binned as white blood cells, since the average cell diameters of CTC’s fall within the 10 - 15µm range.[35–37] A recent review also stated that microfluidic channel size can induce critical flow velocities resulting in shearing and disaggregation of CTC clusters.[18, 38] Another review by Amintas *et al*. concluded that individual CTC isolation techniques are typically not compatible with CTC cluster isolation. They identified three key requirements stating: “Future developments should aim towards methods that allow phenotypic, molecular, and even functional analyses”.[39]

### New Metastatic Cancer Cell Cluster Isolation Methods

New approaches are needed to capture MCCC’s. We have developed a highly selective capture technology with a positive capture bias toward MCCCs by specifically exploiting the surface molecular signature found on most solid tumor cancer cells. Our selective capture approach is based on a biomimetic CD44-induced cell cluster margination effect combined with immobilized capture antibodies coated within the inner walls of a microfluidic chip. The CD44 surface expression is an abundant marker of MCCCs, upregulation of which closely correlates to their metastatic potential.[40–44]

We are reporting on the success of our highly selective capture platform, focused on lung cancer, that preferentially isolates cell clusters predisposed to metastasis. Our technology allows routine and efficient capture of MCCCs providing insights into their biology as well as elucidation of new anti-metastatic therapeutic targets. Ultimately, this will expedite the development of new drugs capable of disrupting clusters and/or cluster formation.

## Materials and Methods

### Microfluidic Chip Manufacturing and Coating Procedure

Design, manufacturing, and coating of the microfluidic chips has been published previously.[45] Production of the microfluidic chips used for this study were outsourced to a commercial partner, µFluidix (Scarborough Ontario, Canada). Each chip was produced from two separate layers of polydimethylsiloxane (PDMS) which were then precision aligned to create our open channel microfluidic chips. The design parameters include 32 individual channels (100µm H x 200µm W each) along with a ‘ceiling’ feature composed on alternating patterns of recessed chevrons, 50µm high which induces chaotic flow pattern.[45]

Ultrapure pharmaceutical grade alginate (Novamatrix; Sandvika, Norway) used for the hydrogel substrate of the Smart-Coating™, was derivatized with streptavidin (AAT Bioquest, Cat.#16885) molecules using carbodiimide chemistry via the carboxylic groups of the sugar residues.[46] Coating of the microfluidic chips with a streptavidin derivatized, calcium (Ca^+2^) cross-linked alginate hydrogel was performed as described previously[45] A secondary coating consisting of a mixture of biotinylated hyaluronic acid and monoclonal antibodies (mAbs) was added to the alginate hydrogel. The secondary coating was prepared using cosmetic grade hyaluronic acid (HA) powder (Resurrection Beauty; Cat.# High-MW-1800kDA) that was biotinylated and Cy5- labeled (1:1:1 molar ratios) using carbodiimide chemistry via the carboxylic groups of the glucuronic acid moiety using Biotin-Hydrazide, Thermo Scientific, Cat.# 21340; and Cy5- Hydrazide from ApexBio, Cat.# A8145. The labeled HA plus a combination of biotinylated monoclonal antibodies was mixed at a 1:16 ratio into phosphate buffered saline (PBS, 1X Caisson Labs PBL01-500ML) along with 0.1% Tween-20.. The 4.5 mL mixture was infused onto the alginate coated chip at 200 uL/min and the chip was subsequently washed with 2.5 mL of PBS at 2.5 mL/min. After the second coating was complete chips were stored in a sealed conical tube containing PBS at room temperature until needed. Figure S1 shows a successful coating of our chip with the fluorescently labeled biotinylated hyaluronic acid-Cy5 (bHA-Cy5) bound to the streptavidin derivatized, cross-linked alginate hydrogel.

Three surface epitopes, EGFR (ERBB1), MET and HER3 (ERBB3) were selected based on published reports that they are typically expressed on the surface of lung cells.[47–49] Three commercially available antibodies that bind to these epitopes were selected: biotinylated anti-EGFR (Santa Cruz Biotechnology, Cat.# SC-120B) biotinylated anti-MET (Cell Signaling Technology, Cat.#64526BC) and biotinylated anti-HER3 (LS Bio Cat.#LS-C87995). All purchased antibodies were tested to confirm cell surface binding on established NSCLC cell lines, HCC827 (ATCC, CRL-2868) and A549 (Angio-Proteomie cAP-0097GFP) by immunofluorescence. Cell line authentication was based provided in vendor information. See Supporting Information.

Experiments to assess the microfluidic chip capture efficiency were conducted with multi-cellular co-cultured spheroids, generated using established cell lines mixed with the non-cancerous human foreskin fibroblasts, HFF cells, as cancer associated fibroblast surrogates. A known number of model NSCLC MCCCs using co-cultured spheroids (1:1 ratio of NSCLC HCC827 and non-cancerous human foreskin fibroblasts, HFF cells) were spiked into normal donor whole blood samples and processed in triplicate. Results comparing zero antibodies, (Smart-Coating™ biotinylated hyaluronic acid / alginate hydrogel only) to 1 antibody (anti-EGFR) and 3 antibodies, (anti-EGFR, anti-MET and anti-HER3 combined) were performed. See the Supporting Information for additional details.

### NSCLC Patient Whole Blood Processing

After IRB approval, a total of eight patient samples (mean age male [3]: 70.9, ssd 1.1; female [5]: 57.1, ssd 1.2) were collected from the University of California, San Diego (UCSD) Moores Cancer Center, an NCI-Designated Comprehensive Cancer Center (UCSD Moores Cancer Center, Project ID# INMC-044, IRB# 181755). Patients previously diagnosed with non-small cell lung cancer (NSCLC), at stages III-A, III-B, III-C, IV-A, IV-B and agreeing to participate in a study, graciously consented to provide whole blood samples. Patient information was compiled for each blood sample, including age/gender, confirmed diagnosis, and cancer stage, at the time of blood draw. Patient’s sex and demographic information was not considered as inclusion or exclusion criteria for this proof-of-concept study. Future studies involving a higher number of patients will consider these factors to help understanding existing disparities. Sample processing was blinded to patient information.

Collection tubes were treated with EDTA and Tirofiban to prevent coagulation and platelet aggregation respectively. Samples were transported on the same day at ambient temperature and processed through our microfluidic system within 8 hours. In an effort to preserve sample integrity to the fullest extent, no red blood cell (RBC) lysis or whole blood pre-processing was performed, since reports of CTC loses have been noted due to pre-treatment methods.[50, 51] The entirety of the patient collected sample was aspirated into a 10mL syringe, attached to a syringe pump and processed through our coated microfluidic chip at a flow rate of 100µL/minute. See Supplementary Methods for additional details.

Flow was not disturbed until the entire sample had been infused through the microfluidic chip. After processing our patient blood sample, the chip was flushed with PBS at a rate of 100µL/minute for 5 minutes to remove any remaining unbound blood or white blood cells. A 4% formaldehyde, methanol-free (PFA) solution was then pumped through at 100µL/minute followed by another PBS wash for 5 minutes. Five of the patients’ samples were processed for immunofluorescence imaging and analysis. Cells were stained with a 10 minute flush of our staining palette consisting of Hoechst 33342 live nuclear stain, (Thermo Scientific Cat.# H3570), anti-CD44-FITC (Abeomic Cat.# 10-7516-F) and anti-EGFR-AF594 (Santa Cruz Biotechnology Cat.# SC-120-AF594) all diluted in PBS, followed by a 5 minute PBS wash, all at the 100µL/minute flow rate.

Our processed chip was imaged across all flow channels using the Yokogawa CQ1 scanner by stitching together 936 fields captured using a 10x objective. Chips were scanned for Brightfield, Hoechst staining (405nm), CD44 (488nm), EGFR (594nm), and our bHA-Cy5 chip coating (647nm). A relative increase of CD44 and EGFR fluorescent signal against background, including well-defined nuclei morphology, were essential gating parameters used to identify potential MCCCs. Using our 10X stitched composite image as a map for locating potential MCCCs, we re-imaged at pre-selected points using a 40x objective to confirm identification. Using a relatively strict inclusion criteria, any irregular nuclei illuminated via the Hoechst signal even if coupled with positive CD44 and EGFR signal were rejected as MCCC candidates. Furthermore, at least 3 individual nuclei had to be discernable in order to be classified as a candidate MCCC.

### NSCLC Whole Blood Processing for RNA-seq Analysis

Three additional patient LC-6, LC-7, LC-8 (selected at random, see Table 1) whole blood samples were processed for RNA-sequencing analysis. Following whole blood infusion, a modified staining protocol was used to reduce imaging time and enhance RNA stability. The chips were flushed with Hoechst 33342 stain (without adding the anti-EGFR and anti-CD44 mAbs), at 100uL/min. for 15 minutes. The Hoechst stain was flushed out using a DMEMs- phenol red media (Thermo Fisher, Cat.# 11965084) at 100uL/min. for 5 minutes..

**Table 1:**
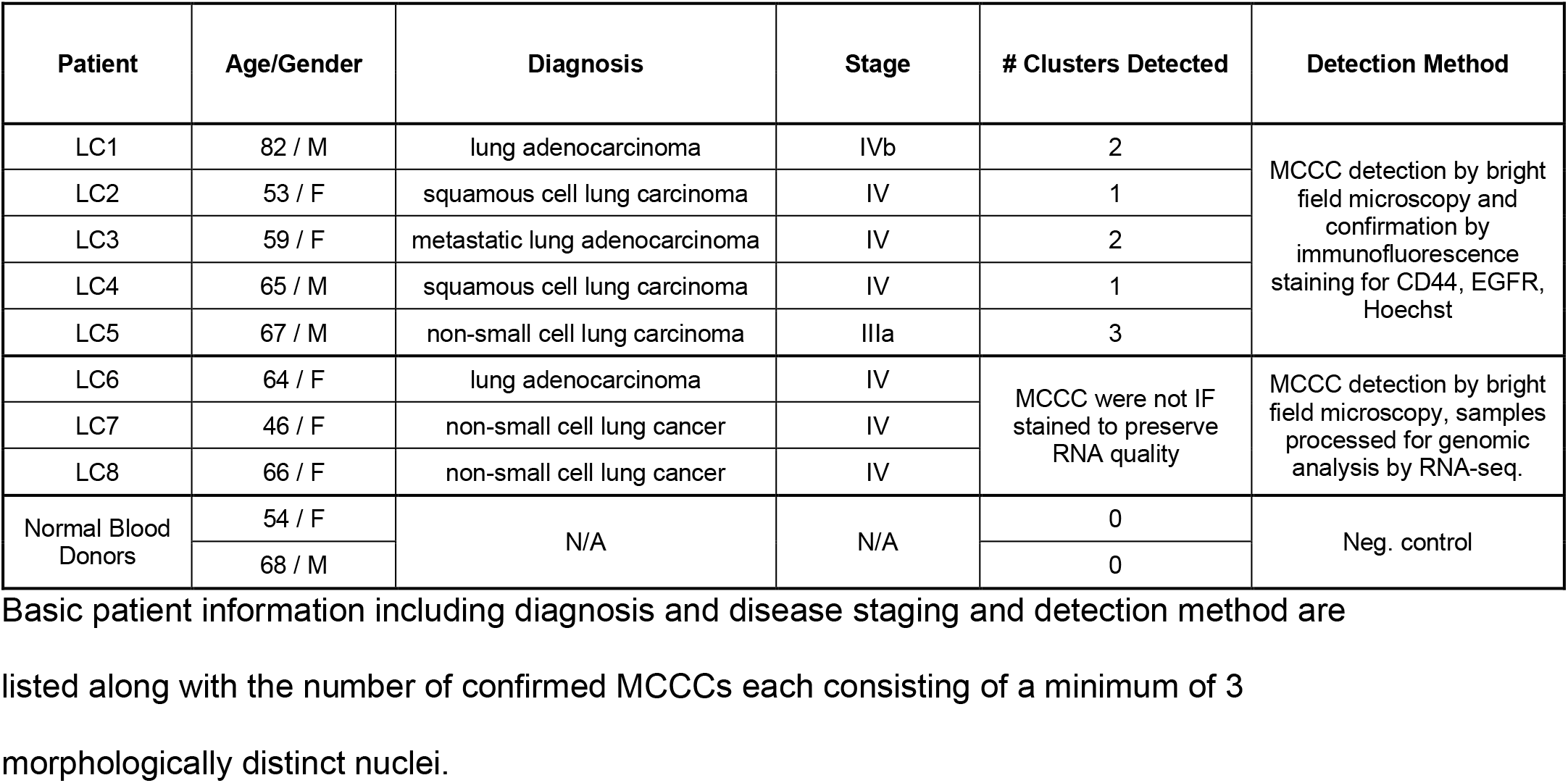
Basic patient information and the number of MCCCs confirmed.

The Inlet and Outlet ports were sealed and imaged using fluorescence microscopy. Brightfield and 405nm images were collected to allow visualization of the captured MCCC nuclei. Post imaging, RNA lysis / stabilization reagent (TaKaRa, Cat. # STO948) was flushed through the microfluidic chip and incubated at zero flow for 15 minutes. Following lysis, fractions were collected at the Outlet port every 15 seconds providing a total of 8 fractions, each with an approximate volume of 25µL. Fractions were immediately frozen at −20°C and sent for RNA sequencing analysis.

One of the patient samples, LC-6 was chosen for RNA-seq analysis. This sample corresponded to a 64-year-old female patient, who was diagnosed by transthoracic biopsy with a TTF-1 positive Stage IV lung adenocarcinoma that had metastasized to the brain. Three RNA lysis aliquots of the LC6 sample were processed and library preparation completed on the 3 aliquots with SMART-Seq® HT Kit (Takara) and Illumina Nextera reagents for tagmentation. RNA sequencing was performed at The Scripps Research Institute NGS core facility. The resulting FASTQ files were downloaded and imported into R. All subsequent analyses were performed using R/Bioconductor packages. A MultiQC v1.7 bioinformatics analysis was generated, revealing the following QC summary data: Reads: 45.1M, % Alignment 94.7% and Phred Quality score ranging from: 31.64 - 33.39.

## Results

### Capture Optimization

Our strategy was to develop a more ‘selective’ technology within a microfluidic system that would preferentially isolate and capture specific metastatic cell clusters rather than utilize a filtration-based or other size-based methodology. The premise was based on the extreme rarity of MCCCs among billions of red and white blood cells thus promoting the need to use an existing molecular biology process to enhance our cell selectivity. Leukocyte extravasation is a natural process involved with tissue damage, inflammation or infection.[52] Within the circulatory system, binding of CD44-presenting cells to endothelial hyaluronic acid (HA) is the first and essential step initiating margination, rolling, and extravasation.[53] Taking advantage of this existing biological principle, we designed and manufactured a microfluidic chip with a Smart-Coating™ technology that combines a tunable biomimetic and immuno-based dual capture approach.[45] An advantage of using our Smart-Coating™ is the ability to vary the number of monoclonal antibodies and biomimetic tethers which are integrated into the coating of the inner wall channels within the microfluidic chip. Immobilized streptavidin was derivatized on the cross-linked alginate hydrogel as previously described.[45] Antibodies with a biotin conjugate are easily bound to the streptavidin within the coating. Combinations of antibodies as well as biotinylated hyaluronic acid, which serves as a biomimetic tether, make up the second layer of our Smart-Coating. Since our platform is tunable, incorporating different biomimetic tethers and monoclonal antibodies enables the system to be adapted to other solid tumor cancer types.

### Capture Antibody Selection for NSCLC

Identifying biomarkers on the surface of lung cells, that are relatively specific for lung tissue, is an important element of our platform. Multiple epitopes were utilized in our immuno-capture method, since this would likely isolate a larger diversity of MCCCs in circulation from different lung cancer patients. Experiments to assess the capture efficiency were conducted using multi-cellular co-cultured spheroids which were counted and spiked into normal whole blood samples. The co-cultured spheroids generated using established NSCLC cell lines resemble the MCCCs found in patients’ whole blood. As shown in Fig. 1, using 3 capture antibodies resulted in a 90% capture efficiency, thus illustrating an advantage using multiple antibodies. Therefore 3 capture antibodies were used for subsequent NSCLC patient samples. The ability to capture co-cultured spheroids using the Smart-Coating alone, without any immobilized antibody, is likely due to the CD44 expression found on the co-cultured spheroids that have an avidity for the hyaluronic acid in our second coating.[54, 55]

**Figure 1:**
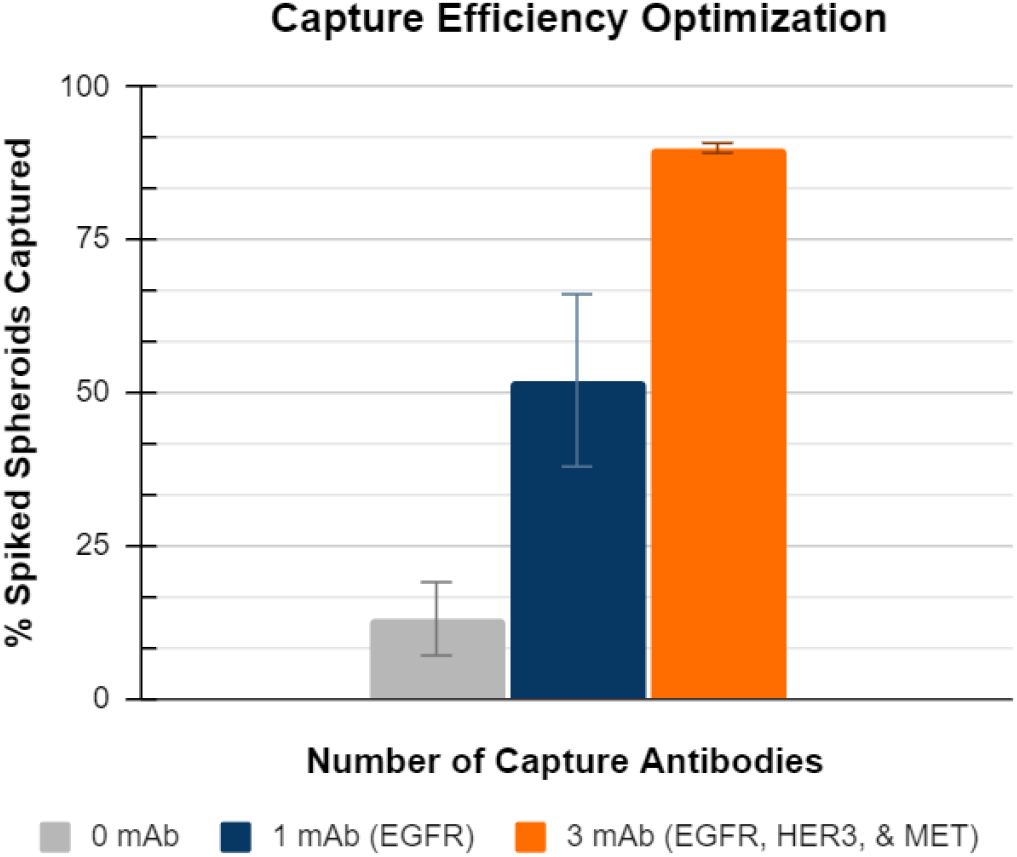
Capture efficiency comparison using varying numbers of tissue specific capture antibodies. The NSCLC co-cultured spheroids were spiked into freshly collected whole blood samples from normal donors. The initial spike count varied for each sample run but consistently fell within the range of 71-91 spheroids. Post processing, NSCLC co-cultured spheroids were quantified via IF. Samples were processed in triplicate.

#### Patient Samples

A total of 8 patient samples were collected from the UCSD Moores Cancer Center and processed through our Smart-Coated™ microfluidic chips that incorporated the 3 monoclonal antibodies (mAbs) which demonstrated the highest capture efficiency. Two additional normal, healthy donor, patient whole blood samples were processed from a 54-year-old female and 68-year-old male, both of which revealed no confirmed MCCCs.

Five of the patient samples, following processing on the chip, included immuno-fluorescence imaging for confirmation of the captured MCCCs using conjugated monoclonal antibodies as listed in Fig. 2. Imaging revealed MCCCs were identified and confirmed in all 5 patient samples, see Table 1.

**Figure 2:**
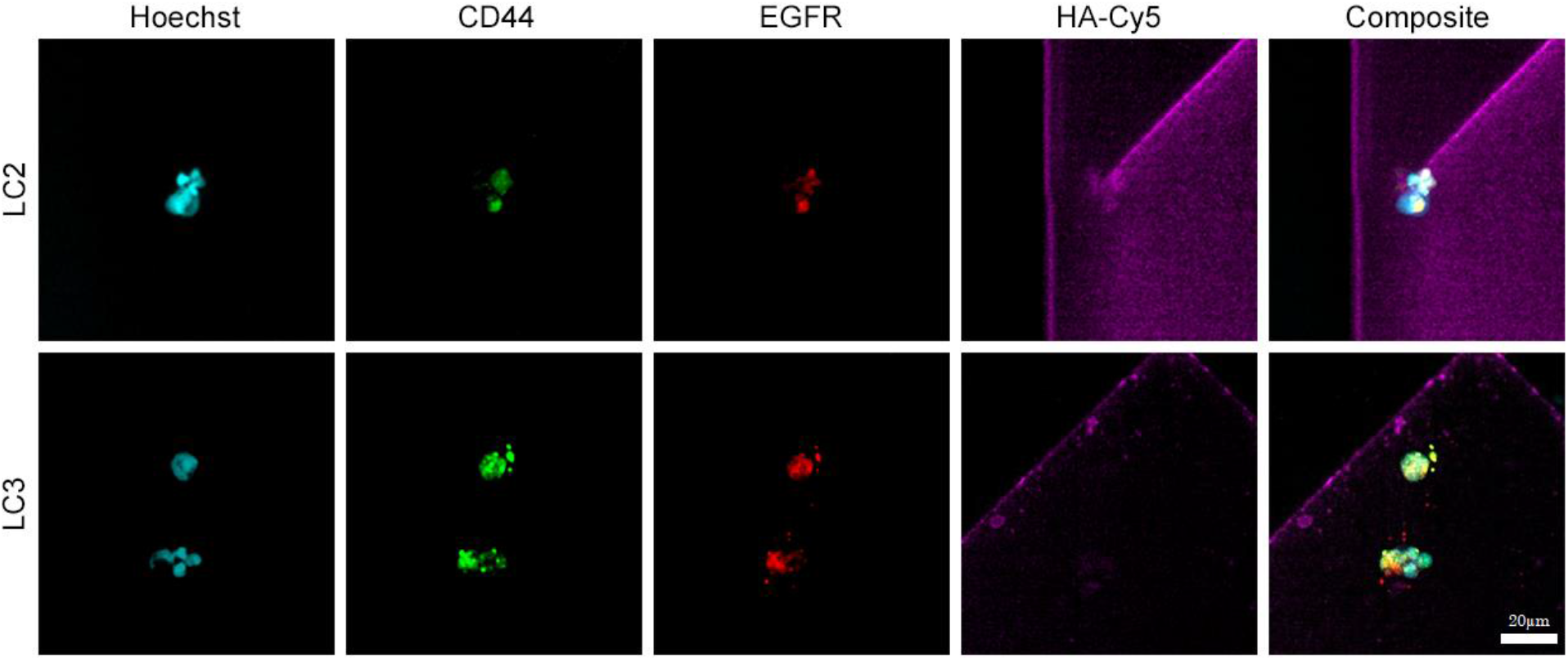
Representative 40X IF images of microfluidic chip captured NSCLC MCCCs from the whole blood of two patients. Nuclei stained with Hoechst 33342 (blue) plus surface antigens anti-CD44-FITC (488nm green) and anti-EGFR-AF594 (594nm red). The Smart-Coating™ is visualized via HA-Cy5 (647nm purple). The overlapping of both CD44 and EGFR fluorescence (yellow) in the composite image indicates cells expressing both markers. Scale bar = 20µm for both LC2 and LC3 images.

Basic patient information including diagnosis and disease staging and detection method are listed along with the number of confirmed MCCCs each consisting of a minimum of 3 morphologically distinct nuclei.

The rationale for the selected surface antigens to confirm MCCCs is that the overexpression of both CD44 and EGFR is associated with cancerous lung cells.[15, 40, 56] Elevated fluorescent signal of anti-CD44 mAb conjugate and anti-EGFR(528) mAb combined with at least 3 distinctnuclei with well-defined nuclear morphology (Hoechst 33342) were essential gating parameters used to identify potential MCCCs. Observations of the captured MCCCs appear to show clusters with multiple nuclei that were relatively compact and not significantly larger than some single cell, non-cancerous monocytes. See Fig. 2.[57, 58] The remaining three NSCLC patient samples were processed with a modified protocol without immunofluorescence to help ensure RNA stability was maintained for subsequent ultra-low concentration bulk RNA sequencing analysis.

Imaged cell clusters were binned into three groups:

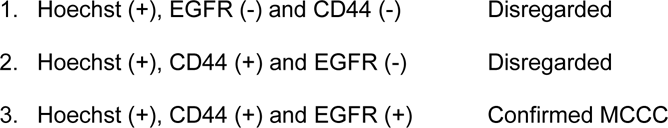

Cells positive for only Hoechst and CD44 were presumed to be white bloods cells.[59] Any clusters with green or red positive staining without clear nuclear morphology via Hoechst staining were disregarded as potential MCCCs. Only clusters with all three stains, including well defined nuclei were considered MCCCs. Examples of the confirmed NSCLC MCCCs captured from whole blood are shown in Fig. 2. Note the heterogeneous shapes and sizes plus the multi-cellular composition as seen by the variation in fluorescence signal distribution among the cells in the MCCCs. A single MCCC was observed in patient LC2 which appears to have 5 nuclei. Also note the apparent compact cellular morphology of the clusters. Two separate, multi-cellular MCCCs were observed in patient LC3. The upper LC3 MCCC appears to have 3 separate nuclei and the lower MCCC has 4 nuclei present.

### MCCC Characterization via RNA-seq Analysis

Determining the unique gene expression profiles represented within the MCCCs via RNA-seq analysis is crucial to learning more about the mechanisms of metastasis. Specifically, the ability to characterize the metastatic cancer cell clusters, captured while in transit, directly from a whole blood specimen will begin to shed light on the numerous signaling pathways involved in the metastatic cascade. Three additional patient samples with confirmed NSCLC diagnoses were processed through our microfluidic chips, using an abbreviated protocol, for the purpose of identifying gene expression profiles represented by the MCCCs. Following a modified processing method, MCCC capture was confirmed with Hoechst staining. After imaging, the microfluidic chip was infused with RNA lysis/stabilization buffer, incubated and then aliquoted fractions were collected at the outlet port.

Understanding the unique gene expression profile emanating from MCCCs found in NSCLC patient’s whole blood is essential to begin discovering new anti-metastatic targets for therapeutic development. The 6^th^ patient, designated LC6, is a 64-year-old female with TTF-1 positive Stage IV lung adenocarcinoma. A 10mL whole blood sample was processed using our Smart-Coating™ microfluidic system. Post analysis imaging revealed multi-nucleated cell clusters (each with 3 or more nuclei based on Hoechst staining). RNA-seq analysis was performed on 3 separate lysate aliquots eluted from the chip processed from patient LC6. Analysis revealed that 138 genes were highly expressed (over 1000 reads) among the MCCCs.

Bioinformatic analysis of the RNA-seq data was used to confirm that the MCCCs captured originated from lung tissue. Expression levels were based on 6 well known genes found in lung cells from previously published datasets of human lung tissue, GSM18949 and GSM18950.[60, 61] The boxplots shown in Fig.3 also included normalized gene expression levels found in lung CTCs and lung primary tumor cells published elsewhere.[61] Our MCCC data was integrated with the GSE74639 dataset. This RNA-seq dataset is comprised of lung tumor circulating single cells, designated (SCs) in the original manuscript, but also referred to here as CTCs as well as lung primary tumor cells (PTs).

**Figure 3:**
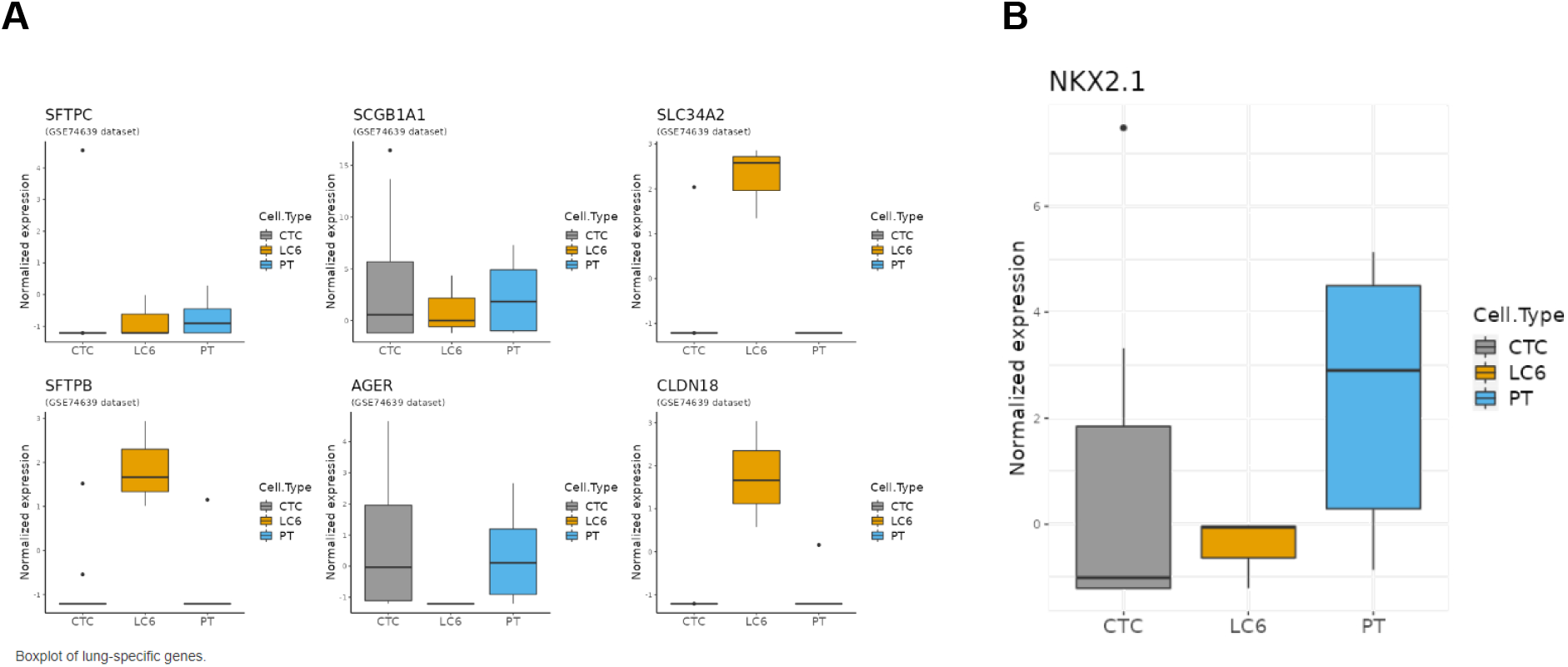
RNA-seq gene expression comparisons between lung circulating tumor cells, primary tumor cells and MCCCs. **A)** Normalized RNA-seq gene expression on a logarithmic y axis of 6 published genetic markers correlated to human lung tissue. Also shown are the comparisons of the MCCCs (patient LC6) to lung individual cells (CTCs) and lung primary tumor cells (PT) for those selected markers. The plots from 5 of the 6 markers selected confirm the lung cell origin of the MCCCs. **B)** Boxplot showing the normalized gene expression of *NKX2.1* (aka *TTF-1*) of the MCCCs compared to both lung CTCs and lung primary tumor samples (PT). The detection of *TTF-1* from lung biopsy specimens is typically used to aid in the diagnosis of NSCLC.

Expression of five out of the six genes assessed were found within the MCCCs from patient LC6: *SFTPC*, *SCGB1A1*, *SLC34A2*, *SFTPB* and *CLDN18* thus confirming the tissue of origin for the captured MCCCs was lung. Another gene was identified within the captured MCCCs, *TTF-1* (aka *NKX2.1*), which is commonly used to aid in the diagnosis of primary pulmonary adenocarcinoma. The *NKX2.1* expression is found in 70-90% of non-mucinous adenocarcinoma subtypes.[62] The boxplot shown in Fig. 3 reveals the expression of *TTF-1*(*NKX2.1*) in the MCCCs from patient LC6, which is consistent with the patient diagnosis obtained from the UCSD Moores Cancer Center, see Table 1. Data supports the finding that the MCCCs are from lung tissue and are adenocarcinoma cells.

An important objective of the RNA-seq analysis of NSCLC patient LC6 was to determine if a unique gene expression profile could be discerned for the captured MCCCs in comparison to circulating lung CTCs and lung primary tumor cells. Additional statistical analysis of the aliquoted samples revealed a unique profile of differentially expressed genes when compared to other individual lung cancer circulating tumor cells (CTCs) and primary tumor (PT) lung biopsy samples. Figure 4 is a Principal Component Analysis (PCA) of the integrated data sets comparing lung CTCs, lung PTs and the 3 MCCC aliquoted samples (LC6). The PCA plot shows a tight grouping of the LC6 aliquots that are significantly different than the PT and CTC lung samples, supporting the postulation that a unique gene expression profile exists for the MCCCs from NSCLC patient LC6.

**Figure 4:**
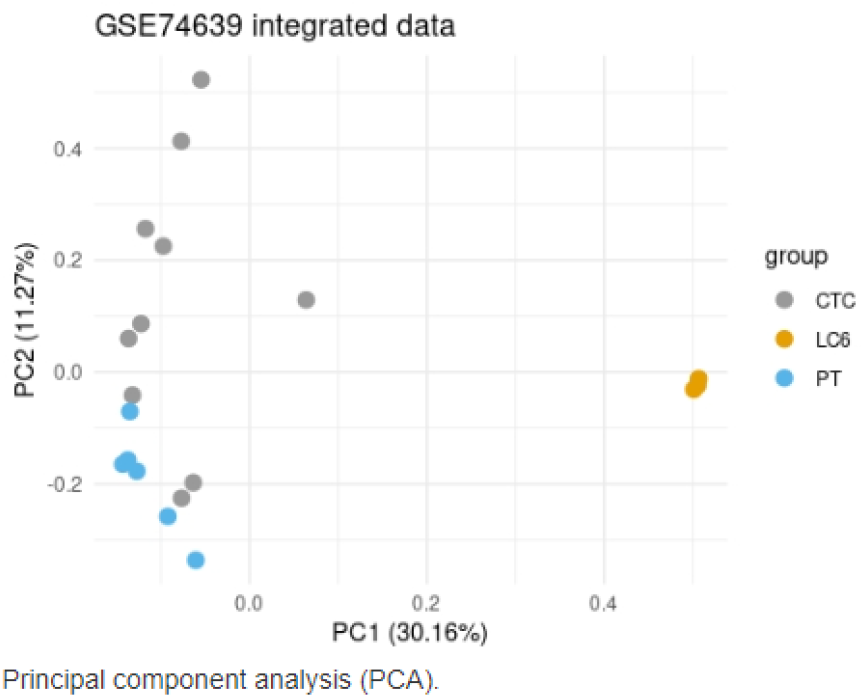
Principal Component Analysis (PCA) of the integrated data sets. PCA comparing lung circulating tumor cells (CTC), lung primary tumor cells (PT) and the 3 MCCC aliquoted samples (LC6). The PCA plot of the patient’s MCCCs sequence data are grouped separately compared to the other lung cancer cell types.

In order to better visualize a unique gene expression profile, based on differentially expressed genes (DEGs) for the MCCCs, a heatmap was generated showing lung MCCCs relative to individual lung CTCs and lung primary tumor cells. The MCCCs from patient LC6 display an intermediate expression profile between lung CTCs and PTs, suggesting some common gene expression among all 3 groups but also showing significant differences as shown in Fig. 5.

**Figure 5:**
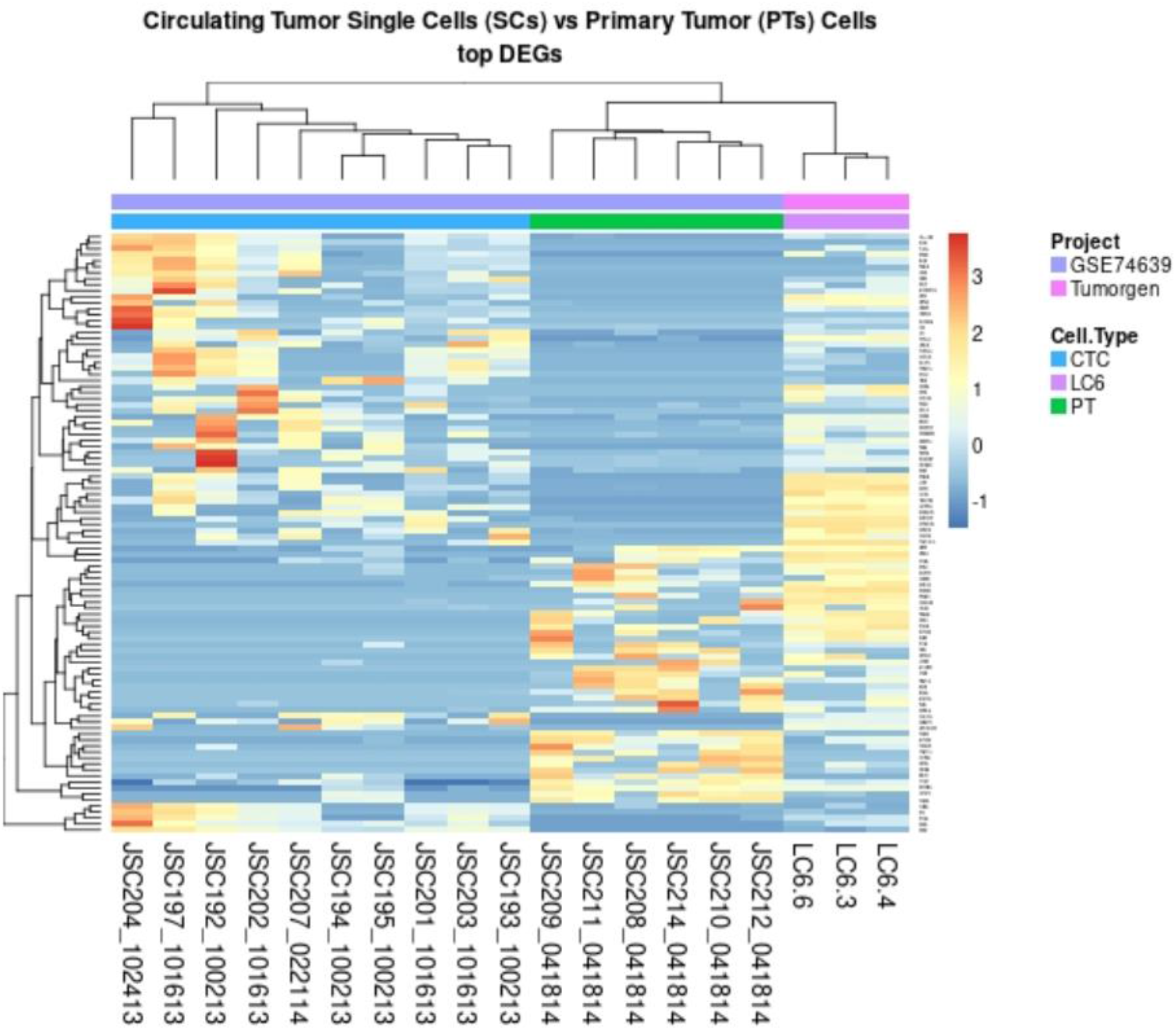
An RNA-seq heatmap was generated using differentially expressed gene (DEGs) between PTs, CTCs and MCCCs. Although the MCCCs share some similarities between lung primary tumor cells PTs and lung CTCs, the DEG grouping of the MCCCs displays a unique profile.

Gene expression mapping of MCCCs suggests a unique profile exists that can reveal pathways driving cancer cell clusters migration to distant metastatic tumor microenvironments.

Using additional bioinformatic data analysis, actionable targets were identified among the MCCCs with corresponding therapeutics available or in development. (Fig.6) Based on the sampling of 9 actionable targets, 3 genes from the MCCCs had noticeably higher expression levels: *EGF*, *ALK* and *ERBB4* (aka *HER4*) compared with lung CTCs and primary tumor cells.

**Figure 6:**
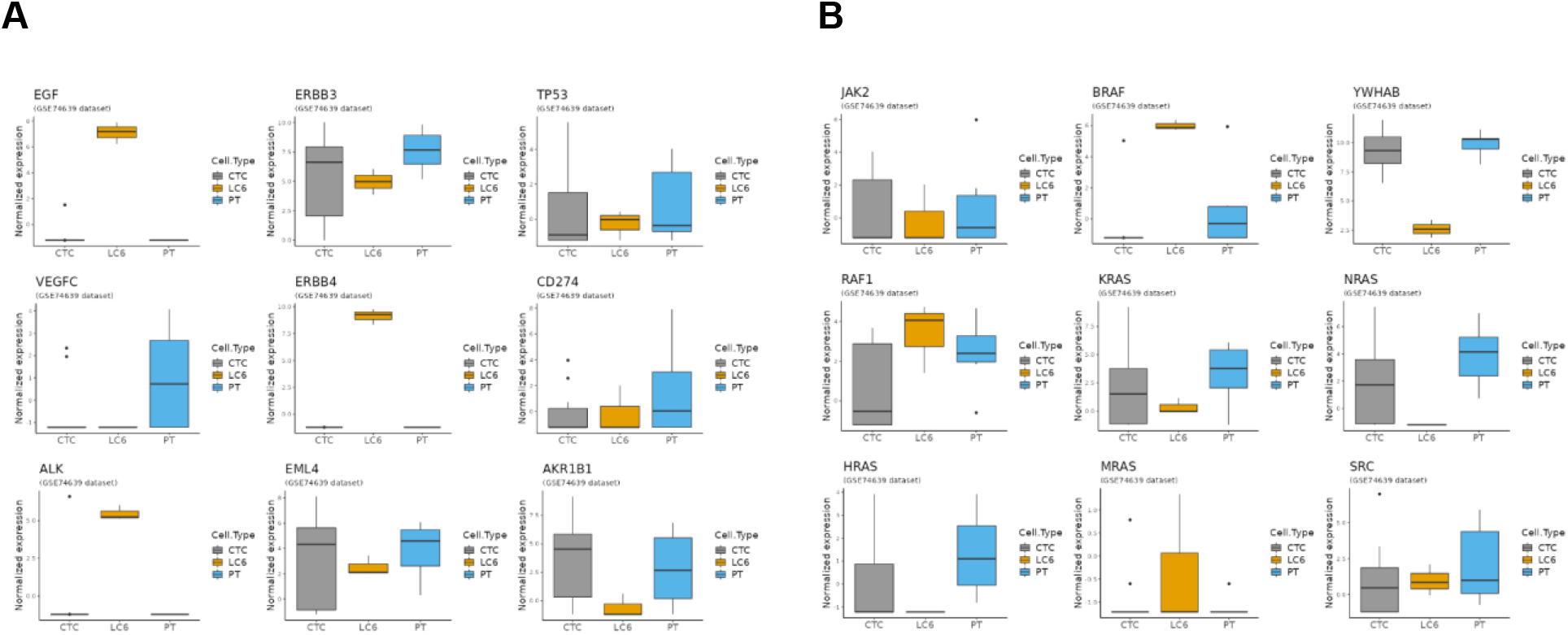
A sampling of actionable therapeutic targets. **A)** Box plots comparing normalized gene expression levels on a logarithmic y axis from a sampling of genes among the MCCCs from patient LC6. The selected actionable genes may be targets since existing drug interventions or new therapeutics are in development. Note the increased expression of *EGF*, *ERBB4* (*HER4*) and *ALK* compared to lung CTCs and lung primary tumor cells. **B)** An Illustration of several genes within the *Ras/Raf* pathway. Note the relatively high expression of *BRAF* among the MCCCs suggesting a potential unseen therapeutic target not seen in lung CTCs or primary tumor cells.

Additional research is necessary to demonstrate that inhibiting the identified targets will interfere with metastatic cluster formation thus ultimately preventing distant metastasis.

Another potential therapy strategy to disrupt MCCCs is to interfere with intercellular interaction and communication. We reviewed the MCCC RNA-seq data to identify elevated expression levels for adhesion and epithelial to mesenchymal (EMT) molecules which may be associated with preparing metastatic clusters for intravasation. Two genes in particular, *COL4A1* and *FGFR1* showed higher expression levels than other lung cell types (See Fig. 7).

**Figure 7:**
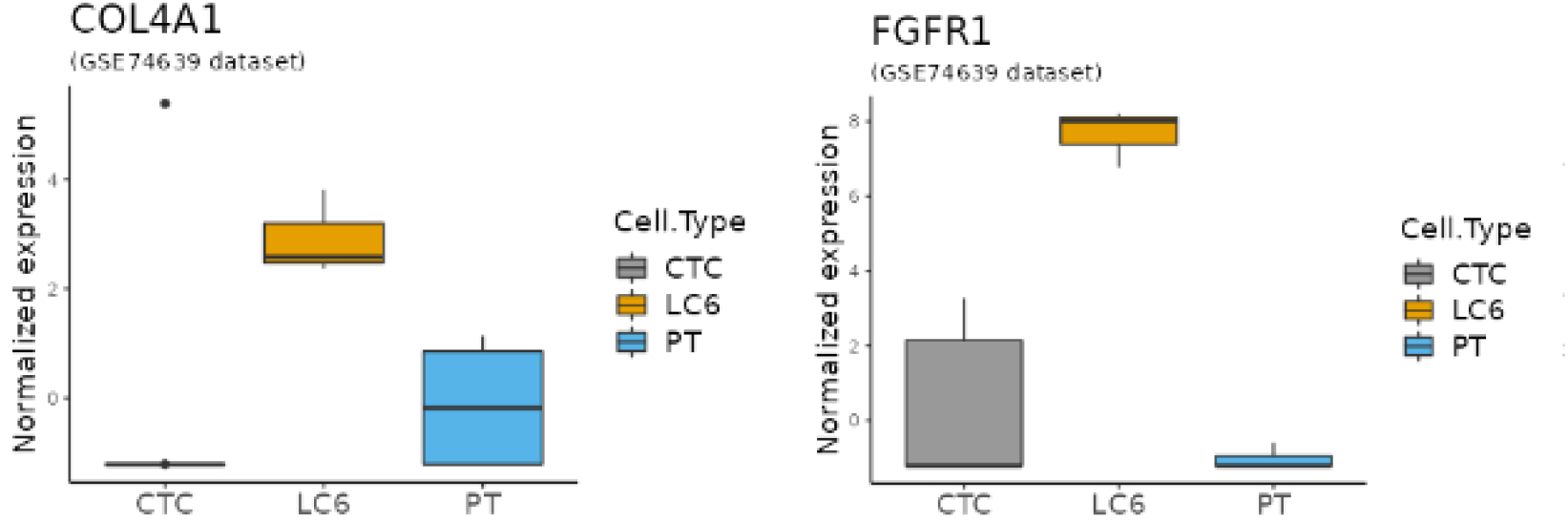
Gene expression correlated to intercellular adhesion and EMT. Two box plots representing *COL4A1* and *FGFR1* normalized expression on a logarithmic scale. MCCCs from patient LC6 appear to show relatively high expression levels compared to individual circulating lung CTCs and lung primary tumor cells.

## Discussion

Understanding the molecular signaling pathways of metastasis is essential for developing new therapeutics designed to prevent cancer’s spread. Isolating and characterizing the cancer cell clusters which are more tumorigenic versus individual cancer cells is an important precursor for new anti-metastatic drug development. [32, 63] Facilitating the isolation and characterization of MCCCs will increase the number of patients analyzed for these metastatic clusters in real time, as well as during various stages of their disease. Technology that allows for frequent sampling at multiple time points may improve our understanding of the metastatic cascade during cancer’s progression. Additionally, easier access to MCCCs will increase research focus on the mechanisms responsible for cancer cluster intravasation and ultimately aid the development of therapeutics to stop their genesis.

Our proof of concept study utilized a completely novel dual capture approach based on a highly selective, biomimetic-immunocapture ‘Smart Coating’ that fast tracks isolation and characterization of extremely rare MCCCs. The Smart Coating™ technology provides a positive selection bias for capturing extremely rare MCCCs directly from unprocessed whole blood. Furthermore, we have shown that a differentially expressed gene profile (DEG) unique to the MCCCs provides evidence that these specific vectors of metastasis require closer observation to better understand the mechanisms responsible for distant metastatic lesions.

The RNA-seq heatmap, which illustrates a clear DEG profile, (Fig.5) further supports our proof-of-concept study, demonstrating the ability to capture MCCCs in circulation and characterize them to identify potentially relevant therapeutic targets. Also, identifying a unique DEG profile for MCCCs supports the assertion that different signaling pathways are in effect during the metastatic cascade.[8, 64]

We acknowledge that our study has limitations. Specifically, the number of patient samples processed using fluorescently labeled mAbs to characterize the MCCCs and the corresponding RNA-seq analyses was relatively low. However, the results demonstrating that one or more MCCCs were found in all patient samples processed are highly promising. Our genomic analysis of MCCCs provides evidence that our dual capture technology can be utilized to reveal previously unseen metastatic pathways. Rather than being conclusive, our goal is to showcase the potential of this type of information and prepare a roadmap for future MCCC studies.

The RNA-seq analyses from our captured MCCCs have shown the expression of several genes that are considered actionable relative to targeted therapies in the treatment of cancer. An example of a potential, previously unseen therapeutic target among the MCCCs is suggested by the elevated *ERBB4* expression as seen in Fig. 6A. The *ERBB4* is a tyrosine kinase receptor and a member of the epidermal growth factor receptor (EGFR) family, also known as HER4. The ERBB4 receptor is activated via ligand binding which subsequently impacts MAPKs and PI3K/AKT pathways.[65] Much research continues to elucidate the role of *ERBB4* mutations and expression among many cancer types but it may be associated with proliferative and migratory abilities of NSCLC.[66] While, *ERBB4* mutations and/or high expression levels observed among NSCLC patients have been identified, their clinical significance remains unclear.[67, 68] Elevated levels of ERBB4 among MCCCs, confirmed when additional patient samples are processed, suggests a possible role in the metastatic cascade thus warranting additional study.

Another gene with high expression levels within the MCCCs relative to lung CTCs and primary tumor is *BRAF* (See Fig.6B). Detection of mutated *BRAF* has been low as reported in only 2-7% of advanced NSCLC patients.[69, 70] The *BRAF* gene is part of the MAPK/ERK signaling pathway, which plays a crucial role in regulating cell growth, differentiation, and survival. Mutated *BRAF* is a well-known driver of various cancers but not usually observed with lung cancer.[71] However, a recent publication identified elevated wild type (WT) *BRAF* expression levels at all stages (I-IV) from lung adenocarcinoma patients and was significantly associated with decreased overall survival.[72] Identifying elevated *BRAF* expression among these rare MCCCs reinforces the need to characterize the metastatic clusters to reveal previously unseen therapeutic targets.

An important strategy for disrupting MCCCs (aka CTC clusters) was described by Nicola Aceto’s group where they identified molecules which interfered with intercellular adhesion.[32] Results from an animal model study using patient derived xenografts demonstrated an 80-fold reduction in metastatic index with the adhesion molecule inhibitor treatment group.

Two genes associated with EMT and adhesion, *COL4A1* and *FGFR1,* showed elevated gene expression (see Fig. 7). The *COL4A1* (Collagen, type IV, alpha 1) is known to promote the growth and metastasis of hepatocellular carcinoma cells.[73] Collagen, a major component of the extracellular matrix (ECM), plays a significant role in cancer metastasis. Cancer cells can alter the composition and organization of the ECM within the tumor microenvironment. Tumor cells often produce enzymes such as matrix metalloproteinases (MMPs) that degrade and remodel the ECM, allowing cancer cells to invade surrounding tissues more easily.[74] A recent publication discusses targeting specific focal adhesion and ECM receptor pathways associated with *COL4A1* for liver and lung metastases.[75]

Also shown is the upregulation of *FGFR1* (Fibroblast growth factor receptor 1) which has been associated with the lung cancer progression pathway in some studies.[76] *FGFR1* is a receptor tyrosine kinase also linked to the *MAPK/ERK* pathway and overexpression of this gene was found to promote epithelial–mesenchymal transition (EMT) plus metastasis.[77] An interesting note regarding elevated expression of *FGFR1* was its link to decreased efficacy of EGFR tyrosine kinase inhibitor (TKI) treatments.[78]

## Conclusion

Understanding the molecular signaling pathways of metastasis is essential for developing new therapeutics designed to prevent cancers’ spread. Isolating and characterizing the metastatic cancer cell clusters, which are more tumorigenic versus individual cancer cells, directly from cancer patients, is essential for finding unseen targets for new anti-metastatic drug development. Expanding research efforts have highlighted the highly promising therapeutic value in targeting and disrupting MCCCs directly in the fight against cancer metastasis.

Identification of *ERBB4*, *BRAF*, *COL4A1* and *FGFR1* as previously unseen therapeutic targets, observed in highly tumorigenic MCCCs, may serve as focal points for new anti-metastatic drug development. Although the gene expression data is based on replicates from a single patient sample, the results demonstrate a proof of concept for our approach. The simplicity and utility of our new platform has the potential to greatly expand the study of metastatic cancer cell clusters for many different solid tumor cancers.

## Author’s Contributions

Conceptualization: Peter Teriete, Jeffrey K. Allen

Formal Analysis: Hector Hernandez-Vargas

Funding Acquisition: Jeffrey K. Allen, Peter Teriete

Investigation: Kourosh Kouhmareh, Anukriti Bhadada, Francisco Downey

Methodology: Darren Finlay

Project Administration: Jeffrey K. Allen

Resources: Jeffrey K. Allen

Supervision: Erika Martin

Writing – Original Draft Prep.: Jeffrey K. Allen, Kourosh Kouhmareh

Writing – Review & Editing: Jeffrey K. Allen, Peter Teriete, Anukriti Bhadada

## Supporting information

https://www.dropbox.com/scl/fi/nevo0gdrxukd79m25mpde/BioRxiv-Supporting-Information-1-Nov-2023.pdf?rlkey=qafma9uj3h5pw8dzkpozhhcu6&dl=0

## Acknowledgments

We are truly grateful for the participation of the patients that consented to provide blood samples for our study. Also, we very much appreciate the support of Dr. Sandip Patel and Dr. Greg Botta from the UCSD Moores Cancer Center. Additionally, thanks go out to Steve Head, Director of the NGS Core facility at Scripps Research Translational Institute for aiding us with the RNA-seq sample preparation and analysis. Further, we appreciate the support provided by Dr. James Evans at PhenoVista Biosciences. The study was supported with an SBIR grant from the National Cancer Institute, 1R43CA261362-01. https://reporter.nih.gov/project-details/10255872

## Note

## Supporting Information

