## Supplementary material for "Capture of circulating metastatic cancer cell clusters from a lung cancer patient can reveal a unique genomic profile and potential anti-metastatic molecular targets: A proof of concept study": https://www.dropbox.com/scl/fi/nevo0gdrxukd79m25mpde/BioRxiv-Supporting-Information-1-Nov-2023.pdf?rlkey=qafma9uj3h5pw8dzkpozhhcu6&dl=0

**Title:**

**Note**

**Supporting Information**

**Supporting Information: Methods**

Our microfluidic system can facilitate the capture of extremely rare MCCCs. It is the first platform specifically designed for this purpose and includes key features, such as channel dimensions suitable for clusters of varying size and structural features introducing non-homogeneous flow to facilitate immuno-based capture. We further addressed rare cell cluster isolation challenges by developing a dual capture mechanism, which combines biomimicry with immuno-capture. Our dual-capture approach is based on a biomimetic cell margination effect driven by CD44 combined with an immobilized antibody capture molecule. The CD44 surface antigen is an abundant marker of MCCCs, upregulation of which closely correlates to their metastatic potential.[1-5] During the normal biological process of leukocyte extravasation into sites of infection or inflammation, the binding of leukocyte presenting CD44 to endothelial hyaluronic acid (HA), its principal ligand, is the first and essential step initiating margination, rolling, and extravasation.[6-8] Tethering of the $CD44^+$ cells by HA allows subsequent interactions, such as PSGL-1 binding to E- or P-selectins as well as VCAM1,[6, 9] to provide full adhesion before diapedesis occurs. The CD44 is further involved in activating signaling pathways required to complete various steps along the active extravasation of the leukocytes.[6, 10, 11] High-jacking of CD44-driven active extravasation further postulates how $CD44^+$ MCCCs increase their metastatic abilities. This means selection of MCCCs based on a $CD44^+$ phenotype confers a bias of the platform toward malignancy, a concept that is supported in a number of recent studies.[4, 12, 13]

The microfluidic chips received from µFluidix underwent a detailed quality control check before coating. A stitched composite of over 900 brightfield images taken with a 10x objective was scanned and compiled for each chip which was then assessed for structural damage and defects.

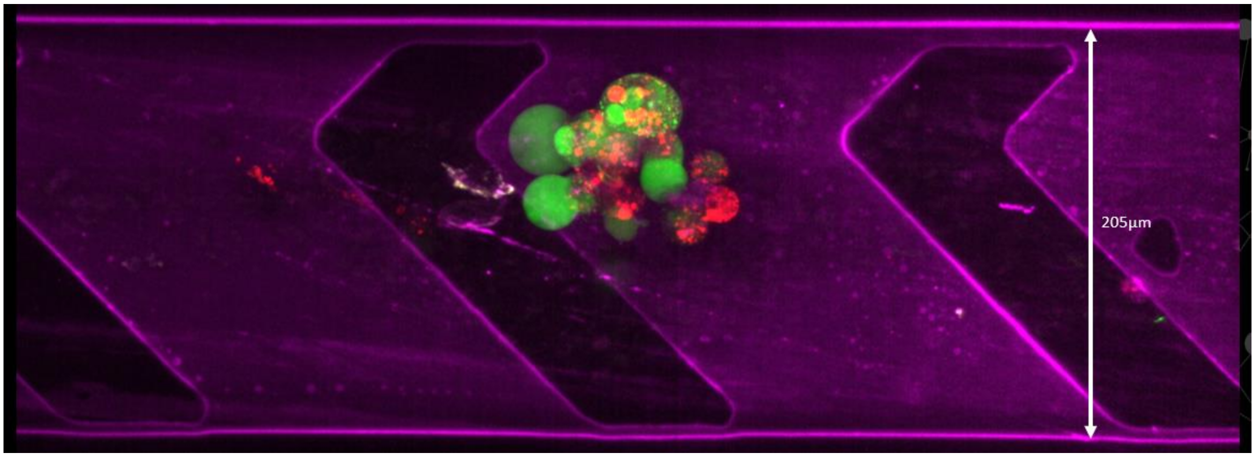

**Figure S1: Immunofluorescence image of a captured model MCCC.** Image of a captured co-cultured spheroid NSCLC A549-GFP (green 488nm) with HFFs (red 568nm) that was spiked into 3 mLs of whole blood and processed through our microfluidic chip. The channel walls shown in (purple 647nm) confirm the presence of our bHA-Cy5 labeled Smart Coating. Maximum intensity projection (MIP) image was acquired using a 40x objective, on a confocal microscope.

Stability testing of our coating has shown functionality is maintained for more than 12 months after coating when stored in aqueous buffer.  Derivatization of hyaluronic acid (HA) (Cosmetic grade, Resurrection Beauty) to incorporate biotin (EZ-Link Hydrazide-LC-biotin, Thermo Scientific, Cat.#21340) functional groups and a Cy5 fluorophore (Cy5-Hydrazide, ApexBio Cat.#A8145) was performed utilizing the same carbodiimide chemistry mentioned earlier.[14].

The capture antibodies used to coat our microfluidic chips were validated to confirm binding to cell surface epitopes using immunofluorescence in both 2D and 3D culture models on our candidate immortalized NSCLC cell lines. All immunofluorescent staining was done on live cells/spheroids to ensure no variations in antibody expression amongst live cells followed by fixation. Cells were seeded in 384-well low volume (Greiner UV-Star 384-well Cell Culture Treated, COC Microplate Cat.788890) and 384-well Sbio U-bottom plates (Sbio MS-9384UZ) and cultured for 72 hours. An antibody mixture containing a 1:100 dilution of anti-EGFR, anti-MET, and anti-HER3 respectively in sterile filtered Goat Blocking reagent (PhenoVista Biosciences proprietary reagent) was added and allowed to incubate in-well for 30 minutes. Cells were then fixed with 4% formaldehyde methanol-free (PFA, Cell Signaling Technology, Cat. # 47746P) for 20 minutes and a corresponding Alexa Fluor secondary antibody was added and allowed to incubate for 30 minutes. (Fixation was necessary to preserve the cells due to lengthy microscope imaging times using a 10X objective to collect over 900 images at 4 wavelengths plus bright field covering all 32 channels in our microfluidic chip). Immediately following processing, both Inlet and Outlet ports were sealed and the microfluidic chip was stored at ambient temperature in PBS until ready for imaging of the captured MCCC's.

Cells were imaged using a fluorescent microscope (Yokogawa CQ1 scanner) and analyzed for positive extracellular specific staining.  Any cells bound to the chip through our immuno-based capture method remain tightly bound since an elevated flow rate of 200uL/min did not remove any bound MCCCs.

Experiments were done using a 3D immortalized NSCLC cell line model system, that were spiked into normal patient whole blood to optimize our Smart-Coating platform. A 1:1 ratio of co-cultured spheroids were generated by seeding NSCLC A549-GFP (Angio-Proteomie, Cat. #: cAP-0097GFP) and/or NSCLC HCC827 (ATCC, CRL-2868) cells respectively, in combination with human foreskin fibroblasts (HFF-1) (ATCC, SCRC-1041) into an 384 well U-bottom, clear, ultra-low attachment plate (Sbio, MS-9384UZ). Cells were incubated at 37°C for 72 hours, resulting in a compact, tightly formed, 30 cell spheroid. Prior to infusion into the microfluidic chip, HCC827 co-cultured spheroids were incubated in-well with CellTracker Green (Invitrogen cat\#\ C2925) for 1 hour until they were ready to be hand-picked, counted and spiked into a 4mL aliquot of freshly collected whole blood. The freshly collected sample from the San Diego Blood Bank was collected, transported and processed on our microfluidic system within 8 hours of donation and kept at ambient temperature. Collection tubes (BD Vacutainer® tube 10mL K2 EDTA, Cat. # BD-367863) were treated with 10.8mg dry coated K2-EDTA and Tirofiban (SelleckChem Cat.# S8594). Injection of either 120 - 200µL of a 5.0mg/mL Tirofiban stock solution in DMSO, 100ug/mL final concentration, was transferred into the 6.0 or 10 mLs of whole blood, depending on the collection tube size used to prevent platelet coagulation engulfing the spheroids or MCCCs. The spiked spheroid blood solution was aspirated into a 10mL injection syringe and attached to a syringe pump. A flow rate of 100µL/minute was used to infuse the blood into our PDMS microfluidic chip. An accurate spiked co-cultured spheroid count was noted and compared to the number of IF imaged spheroids captured on the chip to calculate a capture efficiency percentage. The 100µL/minute flow rate was utilized to ensure spheroid integrity while flowing through the chip. An illustration of the microfluidic injection process is shown in Figure S2. A simplified depiction of the sample processing along with a combined fluorescence / brightfield image of a captured co-cultured spheroid is shown in Figure S3.

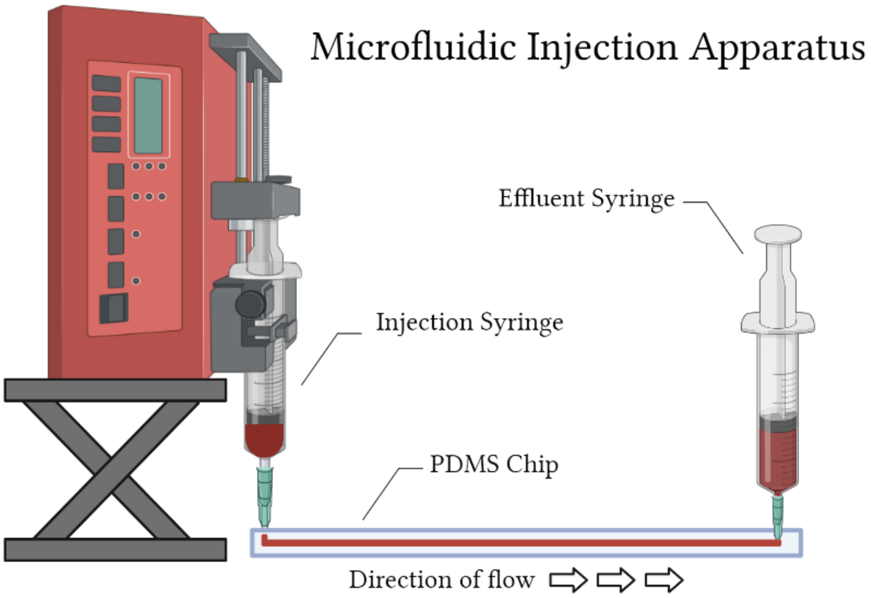

**Figure S2: Schematic of sample processing.** A syringe containing an unprocessed patient blood sample is injected into microfluidic capture chip at a flow rate of 100µL/minute. Effluent can be collected and stored for subsequent multiplexed analyses (ctDNA, exosome, etc.).

**A** **B**

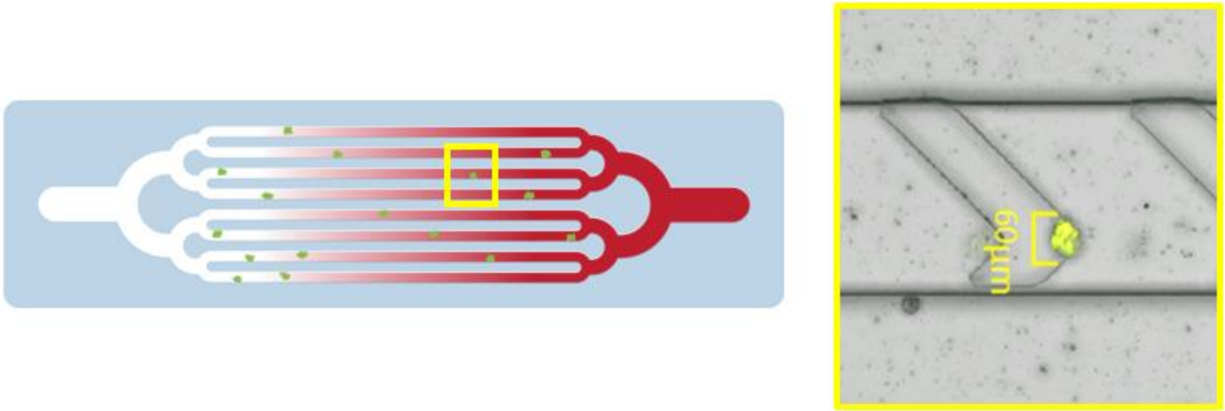

**Figure S3: Simplified illustration of the sample processing and an image of a co-cultured spheroid. A**. An illustration depicting whole blood processing where MCCCs are captured on our Smart-Coating™ within the channel walls of our microfluidic chip, while all remaining RBCs and WBCs are eluted. **B**. Is a composite fluorescent and brightfield microscopy image of one of the captured co-cultured HCC827 spheroid.

**Table S1: Key Resources Table**

| Reagent or Resource | Source | Identifier |
|---|---|---|
| **Capture Antibodies / Coating Reagents** | | |
| Biotinylated anti-EGFR | Santa Cruz Biotechnology | Cat.# SC-120B |
| Biotinylated anti-MET | Cell Signaling Technology | Cat.# 64526BC |
| Biotinylated anti-HER3 | LS Bio | Cat.# LS-C87995 |
| Streptavidin | AAT Bioquest | Cat.# 16885 |
| Sodium Alginate-Pronova-UP-VLVG | NovaMatrix/IFF | Cat.# 4200501 https://novamatrix.biz/store/pronova-up-vlvg/ |
| Hyaluronic acid-comestic grade | Resurrection Beauty | Cat.# High-MW-1800kDa https://stores.resurrectionbeauty.com/hyaluronic-acid-powder |
| **Staining Antibodies / Dyes** | | |
| Anti-CD44-FITC | Abeomics | Cat.# 10-7516-F |
| Anti-EGFR-AF594 | Santa Cruz Biotechnology | Cat.# SC-120-AF594 |

| | | |
|---|---|---|
| Hoechst 33342<br>nuclear stain | Thermo Scientific | Cat.# H3570 |
| **Cell Lines** | | |
| NSCLC - HCC827 | ATCC | Cat.# CRL-2868<br>https://www.atcc.org/products/crl-2868 |
| NSCLC - A549-GFP | Angio-Proteomie | Cat.# cAP-0097GFP<br>https://www.angioproteomie.com/commerce/ccp23644-gfp-human-lung-5bnsclc5d-carcinoma-cells-28a54929-cap-0097gfp.htm |
| Human foreskin<br>fibroblasts-HFF-1 | ATCC | Cat.# SCRC-1041<br>https://www.atcc.org/products/scrc-1041 |

**Supporting Information: Tables**

logcpm_unfiltered.xlsx

logcpm_filtered.xlsx

Excel tables with the normalized data (counts in base 2 logarithmic scale), before and after filtering out of genes with less than 10 counts in average across the 3 replicates.
